# Cytoarchitecture of SARS-CoV-2 infected hamster lungs by X-ray phase contrast tomography: imaging workflow and classification for drug testing

**DOI:** 10.1101/2024.01.21.576083

**Authors:** Jakob Reichmann, Clement Sarrazin, Sebastian Schmale, Claudia Blaurock, Anne Balkema-Buschmann, Bernhard Schmitzer, Tim Salditt

## Abstract

X-ray Phase Contrast Tomography (XPCT) based on wavefield propagation has been established as a high resolution three-dimensional (3D) imaging modality, suitable to reconstruct the intricate structure of soft tissues, and the corresponding pathological alterations. However, for biomedical research, more is needed than 3D visualisation and rendering of the cytoarchitecture in a few selected cases. First, the throughput needs to be increased to cover a statistically relevant number of samples. Second, the cytoarchitecture has to be quantified in terms of morphometric parameters, independent of visual impression. Third, dimensionality reduction and classification are required for identification of effects and interpretation of results. In this work, we present a workflow implemented at a laboratory *μ*CT setup, using semi-automated data acquisition, reconstruction and statistical quantification of lung tissue in an early screen of Covid-19 drug candidates. Different drugs were tested in a hamster model after SARS-CoV-2 infection. To make full use of the recorded high-throughput XPCT data, we then used morphometric parameter determination followed by a dimensionality reduction and classification based on optimal transport. This approach allows efficient discrimination between physiological and pathological lung structure, thereby providing invaluable insights into the pathological progression and partial recovery due to drug treatment.

## 1 Introduction

Tremendous efforts are undertaken to fight infectious diseases such as Covid-19, and to this end constant improvements in scientific methodology are required. One particular recent development of interest is three-dimensional (3D) histophathology based on X-ray phase contrast computed tomography (XPCT) [1–3]. XPCT yields 3D reconstructions of the cyto-architecture with micron-sized or even sub-micron voxel sizes, is compatible with standard tissue preparations such as formalin-fixation and paraffin embedding (FFPE), and effectively adds a third dimension to conventional histology. As computed tomography is intrinsically digital, it comes without any extra step of digitalisation. In fact, it is often even impossible to visually inspect each slice in a stack of several thousands of slices, in particular when it comes to pre-clinical or clinical trials with larger sample size *N*. Digital pathology is no longer an option but becomes a must. To this end, efficient high throughput workflows of automated morphometric analysis and classification are in need. At the same time, data acquisition in XPCT is currently still slow, most investigations remain anecdotic concerning the sample size *N*, and translation from high brilliance synchrotron radiation sources to more accessible laboratory sources is in its infancy.

In the realm of biomedical research, however, the importance of a sufficiently large sample size *N* cannot be overstated, as many confounders may affect the outcome and obscure correlative or causal relationships. At the same time *N* cannot be scaled arbitrarily, in view of animal well-being, ethical requirements, or cost. This can pose challenges, in particular in vaccine or drug development when several compounds have to be tested, and calls for sophisticated and advanced statistical methods. While this generally holds true also for conventional histopathology, the challenges escalate significantly for 3D imaging by XPCT. This is primarily due to the considerable time and human resources required for current image acquisition and data analysis. Workflows are needed which harness the power of automated sample exchange or multi-sample holders, scipts for acquisition, reconstruction and image processing, as well as extraction of quantitative morphometric information.

3D imaging of lung by XPCT is a case in point. The applicability of XPCT has been demonstrated across a spectrum of scales, ranging from the macroscopic to the microscopic [4–13], and even including in-vivo lung imaging. With its intricate 3D networks of airways and vasculature, alveolar ducts, spaces and septae, 3D imaging is desirable and at the same time facilitated by strong contrast based on the density contrast of tissue and empty space filled by air or the embedding medium. XPCT has also been used to image unstained FFPE lung tissue from patients who succumbed to COVID-19, offering 3D insights into diffuse alveolar damage (DAD), hyaline membrane formation, lymphocyte infiltration, vascular damage, and intussusceptive angiogenesis [14].

With the 3D reconstructions at hand, morphometric analysis of the tissue can be carried out. Parameters such as tissue density, surface areas and curvature, sphericity of objects, as well as compactness can be determined, see for example [15]. For lung tissue, sizes of alveoli and thickness of septae are of particular interest [16]. More generally, extracting quantitative image information has become a major effort in view of diagnostic and prognostic capabilities, and in clinical context is often referred to as “radiomics” [17, 18].

Once morphometric features have been extracted, statistical evaluation and classification can be performed. Here, the challenge lies in the high dimensionality. In each sample image (patient, animal), there are typically many instances of the sought-after features. This holds especially true for bulky 3d images. Therefore not individual features, but instead whole collections of features must be compared between samples. These can be interpreted as being drawn from an underlying ground-truth probability distribution function (PDF) that captures the specific properties of each sample. Fortunately, machine learning and in particular optimal transport (OT) has recently evolved as a major tool for quantitative comparison of PDFs. OT provides a mathematical framework of re-arranging ‘mass’ from one location in a PDF to another, while minimizing the global (or average) cost of transport. Since its mathematical formalization by Kantorovich, OT has evolved into a well-known, versatile tool. Due to the availability of increasingly efficient numerical methods [19], several important applications in machine learning and image analysis have emerged, such as image registration [20, 21], segmentation [22–24], pattern recognition [25] and data fusion [26].

In this work, we investigate the effectiveness of several (blinded) drug compounds for treatment of Covid-19 in a small animal model, based on 3D histopathology of lung as the predominantly afflicted organ. Owing to the difficulties in SARS-CoV-2 infection in mice, we turn to the well established hamster model, as introduced in [27, 28]. The scope of the work is primarily in method development, and demonstrating the potential of high throughput laboratory XPCT in combination with automated image processing and statistical analysis based on OT. After exploring different imaging configurations both at synchrotron and in-house X-ray sources, we scanned FFPE tissues of more than 50 hamsters with the same compact *μ*CT configuration and use the so-called chord length distribution as a distinct morphometric measure for the alveolar spaces in hamster lungs. Apart from positive and negative controls, five different drug candidates for Covid-19 treatment are included in the study, and are discriminated based on OT analysis. Interestingly, promising candidates are identified, notwithstanding the still very small *N* and the confounders intrinsic in such a trial.

The manuscript is organized as follows: following this introduction, methods of sample preparation and image acquisition are detailed, as well as the analysis workflow including the concept of chord lengths distribution (CLD), the rudimentary principles of optimal transport, and data processing methods. Subsequently, the results are presented for XPCT image quality, morphometric measures, and OT-analysis. The classification is then applied to test samples of infected SARS-CoV-2 hamster lungs treated with five different drug candidates. The last section combines discussion, brief conclusions, and outlook.

## 2 Methods

### Animals

Male Golden Syrian hamsters (Mesocricetus auratus; RjHan:AURA; 80–100 g) were obtained from Janvier Labs (Saint Berthevin, France) and were housed in standard rodent IVCs Type III in groups of 3 to 4 animals under standardized conditions (22°C; 12/12h light cycle). As diet, rodent pellets and water ad libitum were fed. For acclimatization, rodents were housed under these conditions for one week prior to inoculation. The experiments were conducted in a BSL-3 animal facility. Animals were infected orotracheally with 1×10^5^ TCID_50_ SARS-CoV-2 Germany/BavPat1/2020 (BavPat1) [29] (GISAID accession EPI_ISL_406862) in a volume of 100 μL. The animals’ well-being and body weight were checked daily. Animals were clinically observed for 7 days with a daily sampling for virological analysis. After 7 days, animals were euthanized by inhalation of an isoflurane overdose followed by intracardial exsanguination and decapitation. Virological analysis of the collected samples was performed as described in [28].

### Sample Preparation

In total, 57 hamster lungs divided into nine groups were examined in this study, including three groups of negative, one group of positive controls (*POS* − *CT RL*), as well as five groups of drug treated cases, i.e. hamsters infected and subsequently treated by five different drugs. These are referred to as the five drug groups (drug groups #1 − 5). The three negative control (*NEG* − *CTRL*) groups consist of untreated and uninfected control (*UNI* − *CTRL*), and uninfected control groups subjected to treatment with a drug vehicle but no drug loaded. Corresponding to the solution injected, these are referred to as polyethylene glycol control (*PEG* − *CTRL*), and phosphate-buffered saline (*PBS* − *CTRL*). The positive control group was infected with 1×10^5^ tissue culture infectious doses 50 (TCID50) SARS-CoV-2, but not treated. Both positive and negative controls are used as training data. For clarity, the control samples were further tabulated in Tab.1 and the preparation procedure is elucidated in Figure 1**a**. The five different drug candidates are tested with respect to the metric space spanned by the morphometric features of the negative and positive control groups. The five drug groups are referred to as test data. Additionally, a ‘lung affection score’ (LAS) ranging from zero (healthy) to one (sick), was assigned to every dissected lung by visual inspection.

**Table 1.** List of all hamsters in the groups forming the negative and positive controls, serving as training data. Positive controls are denoted as *POS* − *CTRL*, the negative controls consist of three groups, the uninfected control (*UNI* − *CTRL*), the polyethylene glycol control (*PEG* − *CTRL*), and and phosphate-buffered saline control group (*PBS* − *CTRL*). The macroscopically assessed lung affection score (LAS) is also given.

| Hamster | Group | LAS [%] |
| --- | --- | --- |
| 52 | UNI-CTRL | 0 |
| 53 | UNI-CTRL | 0 |
| 55 | POS-CTRL | 20 |
| 56 | POS-CTRL | 30 |
| 57 | POS-CTRL | 100 |
| 58 | POS-CTRL | 70 |
| 59 | POS-CTRL | 100 |
| 60 | POS-CTRL | 80 |
| 62 | POS-CTRL | 90 |
| 63 | PEG-CTRL | 0 |
| 64 | PEG-CTRL | 0 |
| 82 | PBS-CTRL | 5 |
| 83 | PBS-CTRL | 0 |
| 84 | PBS-CTRL | 0 |

**Figure 1.**
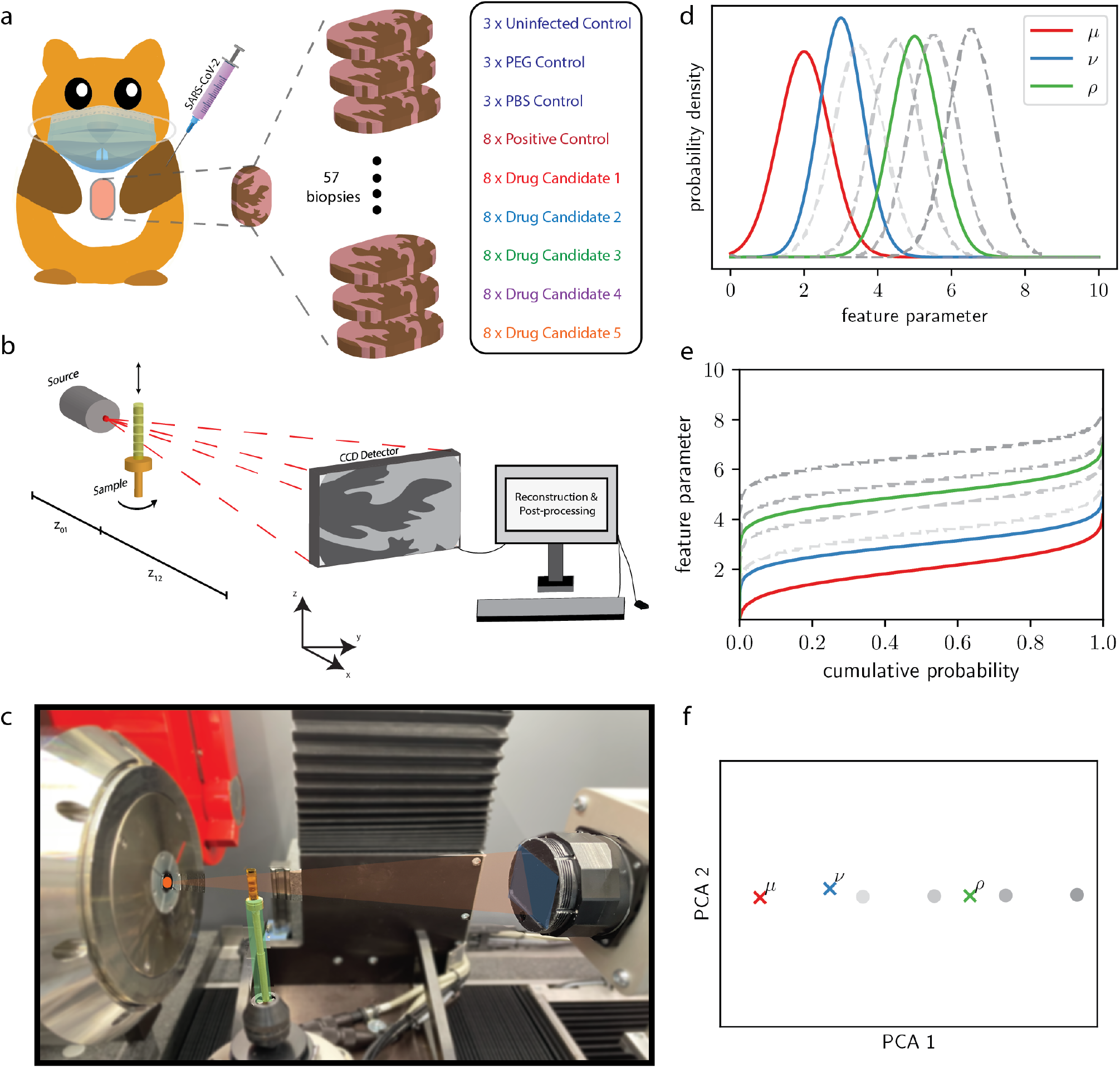
**a** Seven days after infection of Golden Syrian hamsters by SARS-CoV-2, the animals were sacrificed, lung autopsies were extracted and paraffin embedding was performed. In total, 57 hamsters, divided in nine groups, were examined in this study, including uninfected controls (×3), positive controls (×8), PEG (×3) and PBS controls (×3) as well as five different drug candidates (×8, each). In further analysis, uninfected, PEG and PBS controls are referred to as NEG-CTRL, while positive controls are referred to as POS-CTRL. **b** Schematic depiction of the in-house experimental setup. The cone beam geometry allows for an adjustable magnification and thus effective voxel size depending on the distances x_01_ and x_12_. A scintillator-based CCD-detector is recording the incoming photons, followed by a phase- and tomographic reconstruction from the acquired projections. **c** Photo of the experimental setup with the source (left), the sample stage (center, green) and the detector (right, blue). X-rays are created at the anode (orange spot) and propagate through the object, reaching the scintillator where X-rays are converted to visible light and subsequently to an electrical signal. **d,e,f** *From top to bottom, in color:* three synthetic sample distributions *μ, ν* and *ρ*; the corresponding inverse cumulative distribution functions used in (2); the corresponding PCA embedding, obtained by projecting onto the two first principal components. *From bottom to top, in gray:* Some selected points around *ρ* in the 2d-embedding, along the first principal component; approximately corresponding to vertical translation in the inverse CDF space; approximately corresponding to a translation on *ρ* (with a slight change in standard deviation), thus making this direction in the embedding interpretable.

All hamsters were dissected 7 days after infection at the Friedrich-Loeffler Institute (FLI). After lung extraction, the samples were placed in a 10% formaldehyde solution. In a subsequent step, embedding of the tissue had to be performed to ensure stabilization and preservation of the tissue. FFPE is the most common embedding and preservation procedure in clinical pathology, and has already been successfully used for 3D virtual histology of lung with synchrotron and laboratory radiation [14, 30]. For FFPE, the tissue is first dehydrated by immersion in a series of ascending ethanol solutions. Xylene is then used as an intermediate solvent to allow for subsequent wax infiltration, which is not solvable in ethanol. While being placed on a cassette, the tissue is then infiltrated with paraffin.

After solidification, 3 mm biopsy punches are extracted from the identified regions of interest, guided by parallel histological sections. The paraffin embedded tissue biopsies were then mounted on a brass pin for tomographic recording. The tissue fixation, embedding and mounting of samples followed well established protocols described in [1].

### Experimental Setup

Tomographic data was primarily acquired with a commercially available laboratory CT system (EasyTom, RX Solutions, Chavanod), motivated by the fact that such instruments can easily be made available in a pre-clinical setting. Further, they can be used over long times in rather simple and time-stationary conditions, ideal for biomedical studies with large sample numbers *N*. For the scans, the open transmission tube (Hamamatsu) of the EasyTom instrument was selected, equipped with a LaB6 cathode and a tungsten (W) target, and operated at a tube voltage of 60 kV and a target power of 6.6 W. The spot size was approximately 1.5 *μ*m. Projection images were acquired by a CCD detector of 9 *μ*m pixel size (2x2 binned) (Ximea, Münster) with a fibre-coupled Gadox scintillator. Experimental parameters (geometry, acquisition time, source settings) were optimized by comparing the reconstruction quality for selected test samples, and then fixed for the entire series to the values tabulated in Tab. 2. To reduce image noise, four images were recorded at each angle and averaged (median). A Fourier-filter based scheme was used to account for phase contrast. Only in-house data recorded under identical settings was further used for statistical classification (illustrated in Figure 1**b,c**).

**Table 2.** Data acquisition and detection parameters, including the source setting, the source-to-sample distance *x*_01_, the sample-to-detector distance *x*_12_, the resulting effective pixel size *px*_*e f f*_ and field-of-view *FOV* (horizontal and vertical), the number of projections per scan #*pro j s*, as well as the exposure time *τ* and total scan time.

|  | Laboratory | P10 | ID16a |
| --- | --- | --- | --- |
| Tube voltage / energy | 60 kV | 13.8 keV | 17.1 keV |
| $x_{01}$ | 9.3 mm | 88 m | 34.7 mm |
| $x_{12}$ | 93.1 mm | 29 mm | 1.17 m |
| $px_{eff}(\mu\text{m})$ | 1.63 | 0.65 | 0.09 |
| FOV (h × v) (mm<sup>2</sup>) | $3.3 \times 2.2$ | $1.6 \times 1.4$ | $0.29 \times 0.29$ |
| $\#projs$ | 1568 | 3000 | 2000 |
| $\tau$ (s) | 1.2 | 0.035 | 0.2 |
| Tot. scan time (min) | 125 | 1.25 | 180 |

Synchrotron data was recorded for selected samples, serving as a reference (ground truth). Specifically, we recorded XPCT scans at two synchrotron instruments, the P10 beamline at DESY (PETRA III, Hamburg) for microscale XPCT with parallel beam tomography, and the ID16a nano-imaging beamline (ESRF, Grenoble) for nanoscale XPCT (holo-tomography). At P10 a photon energy *E* _*ph*_ = 13.8keV was defined by a Si(111) channel-cut monochromator. A field-of-view (FOV) of approximately 1.5 mm was covered. 3000 projection angles were recorded over a 360° interval, with continuous rotation and a rotational speed resulting in an illumination time *τ* = 0.035s per projection. The entire recording with additional empty beam references before and after the tomographic scan took less than 2 minutes. The detector was positioned at a (defocus) distance of *z*_12_ = 29 mm behind the sample to achieve phase contrast in the direct contrast regime. Images were recorded with a pco.edge 5.5 sCMOS camera (PCO, Germany), equipped with a rolling shutter and a fast scan mode, and achieving a maximum frame rate of 100 Hz. This camera was coupled to a 50 mm-thick LuAG:Ce scintillation screen using a high-resolution optical detection system (Optique Peter, France), equipped with a 10x magnification microscope objective. This configuration resulted in an effective pixel dimension of *px*_*e f f*_ = 0.65*μ*m. The detection and acquisition scheme are described in detail in [31]. Phase retrieval was performed on the projections using the contrast-transfer-function (CTF) approach [32] (single distance), implemented numerically in a Matlab phase retrieval package (HolotomoToolbox [33]).

Since the FOV and pixel size are of of the same order of magnitude, the P10 data can be regarded as a ground truth reference for the study based on inhouse recording, showing the cytoarchitecture at the same scale, but with sharper and more contrasted reconstructions, as well as tractable gray values owing to the monochromaticity and better justified phase retrieval filters. Finally, as a comparison and high-resolution benchmark, nano-holography scans were recorded for two samples, one *POS* − *CTRL* and one *UNI* − *CTRL* sample, using the nano-focusing optics at the ID16a beamline of the European synchrotron radiation facility (ESRF, Grenoble). Taking into account the strongly holographic regime, four distances were recorded for phase retrieval, implemented as a generalized Pagainin method with subsequent iterative refinement [34]). All relevant experimental parameters are tabulated in Table 2.

For all instruments, the acquired projections were saved in either the .tiff or .h5 data format. Tomographic reconstruction was then performed by filtered back projection (FBP) and the Feldkamp-Davis-Kress (FDK) algorithm, for the parallel beam data (P10) and the cone beam recordings (ID16a,EasyTom), respectively. Automatic rotation axis and drift corrections, as well as ring removal techniques were used, where appropriate. At EasyTom, the reconstruction software provided with the instrument was used. P10 and ID16a data was reconstructed using the ASTRA-Toolbox [35] and PyHST [36], respectively. Final reconstructions were stored in the .raw file format. Representative imaging results for all three instrumental settings are shown in Fig.2 of the results section below.

### Chord Length Distribution

In order to quantify changes of the peripheral lung structure associated with SARS-CoV-2 infection of the hamsters, we computed the so-called chord length distribution (CLD) as a characteristic measure [37]. The CLD is well suited to quantify changes in size of the alveolar lumen and septae associated with different pathologies. To this end, the reconstructed gray values representing electron density *ρ*_*x,y,z*_ are binarized into two phases, 0 (lumen) and 1 (septae), using Otsu’s thresholding [38]. For the binarized volume masks, chords are computed. A chord is defined as a segment of a line which traverses the volume with random orientation and intercept. The line is then divided into several segments of length *L*_*c*_, defined by endpoints at the interfaces between the two phases. The procedure is visually explained in Figure 3. For the numerical computation of the CLD, we used Bresenham’s line algorithm [39], implemented in [40]. Given proper normalisation, the CLD represents the probability density of finding a chord with length *L*_*c*_ for each of the two phases of the binarized image. As a quantitative measure for two phase materials, the CLD is well established not only in material science [41–43], but also for (binarized) lung morphology [44]. However, up to now, it was solely used for two-dimensional (2D) histological tissue slices, while we here extend the method to 3D image analysis of lung tissue.

### Optimal Transport (OT)

For an approachable introduction to the topic of optimal transport with a focus on numerical methods we refer to [19]. A more mathematical exposition can be found in [45]. Here we very briefly sketch the central concepts used in this article.

Let *μ* and *ν* be two given probability distributions on ℝ^*d*^ (representing, for instance, two chord length distributions). For transporting *μ* onto *ν*, we introduce a probability distribution *π* on ℝ^*d*^ × ℝ^*d*^, where intuitively *π*(*x, y*) represents the mass density that is taken from *μ* at *x* to *ν* at *y*. For *π* to describe a valid transport plan between the two measures, its first and second marginals must be equal to *μ* and *ν* respectively. We denote the set of transport plans between *μ* and *ν* by Π (*μ, ν*). Let now *c* : ℝ^*d*^ × ℝ^*d*^ →ℝ be a cost function, where *c*(*x, y*) specifies the cost of taking one unit of mass from *x* to *y*. Then the total cost associated with a plan *π* is given by 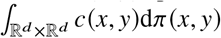. The optimal transport problem then consists of finding the transport plan with lowest cost, i.e.

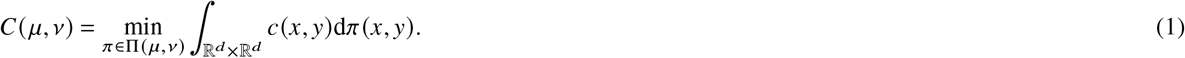

For the choice *c*(*x, y*) = ∥*x* − *y*∥^2^, this defines the 2-Wasserstein distance 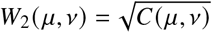 on the set of probability distributions. This distance is particularly relevant for data analysis, since it quantifies geometric discrepancies between data distributions more meaningfully than other common tools, such as the 𝕃^2^-norm (also known as mean squared error) or the Kullback–Leibler divergence (also known as relative entropy).

For one-dimensional distributions, *d* = 1 (such as for chord length distributions), *W*_2_ can be written as 𝕃^2^-norm on the space of inverse cumulative distribution functions (CDF), namely

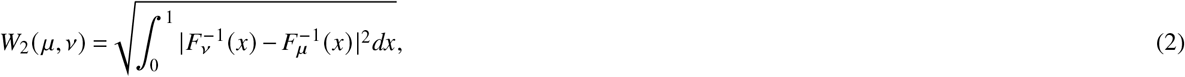

where *F*_*μ*_ is the CDF of *μ* and 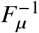 denotes the corresponding inverse CDF. This means that we isometrically embed the samples into the Hilbert space 𝕃^2^ 0, 1. Note that computation of CDFs, their inversion, and evaluation of (2) are numerically simple. For inversion we use one-dimensional interpolation [46]. Standard techniques of statistical analysis in Hilbert spaces can then be applied to the inverse CDF 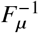 representing our data distributions. For instance, principal component analysis (PCA) can be used for dimensionality reduction and to identify prototypical directions of variation in the set of samples. By projection a sample onto the first few principal components, a low-dimensional embedding of the data can be generated. Conversely, any point in the low-dimensional embedding corresponds to some hypothetical inverse CDF 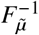 for some hypothetical distribution 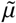. 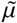 can be obtained by transforming back from the inverse CDF to the original PDF. In this way it is possible to visualize the dominant variations in the collection of samples and to evaluate whether they may be associated with pathological changes. Figure 1**d-e** illustrates, for simple synthetic data, the embedding onto the two dominant principal components, as well as the possibility to compute the inverse of the embedding in order to visualize the transformations that the principal components encode on the PDFs. A thorough introduction to optimal transport in one dimension, including details such as how to handle measures with atoms (i.e. mass concentrated on a single point), is given in [45, Section 2], a tutorial for applications in image analysis can be found in [47]. For our numerical analysis we use the LinOT library for Python.^1^ An approximate generalization of this framework to distributions in higher dimensions has been proposed in [48], see [14] for an application in digital histology.

## 3 Results

In the following we present the results with regard to the three major aims of this study, namely (a) to implement an efficient XPCT imaging workflow for pre-clinical studies, (b) to quantify morphology of lung tissue, and (c) to use OT analysis to test potential drugs in pre-clinical small animal studies of Covid-19.

### XPCT: image quality and workflow

Figure 2 demonstrates the image quality achieved at the laboratory setup in comparison to the synchrotron data, at P10 and ID16a, respectively. As can be seen, the inhouse image quality is sufficient to resolve the general tissue morphology in terms of alveolar spaces and lumen. A differentiation between infected and control tissue seems plausible in terms of the expected thickening of septae. Importantly, the geometric distribution of lung tissue can be captured, or more precisely the morphology in the two phase model, which accounts for air (paraffin) and tissue, respectively. With the parallel beam synchrotron modality, more details of the septae become visible, such as the more strongly contrasted cellular nuclei, while maintaining a comparably large FOV. Finally, high-resolution scans at the ID16a beamline reveal much sharper tissue boundaries, different cell types based on different grey values, as well as sub-cellular structures. This opens up a potential for quantification and classification of pathologies based on cellular and sub-cellular structural parameters (features) [14, 49]. However, it is more challenging to achieve larger FOVs for a representative sub-volume, and more importantly, the required number of animals to be tested. Contrarily, the inhouse configuration combines sufficient data quality for the current purpose with long-time availability, and accessibility, and is hence selected for the current study.

**Figure 2.**
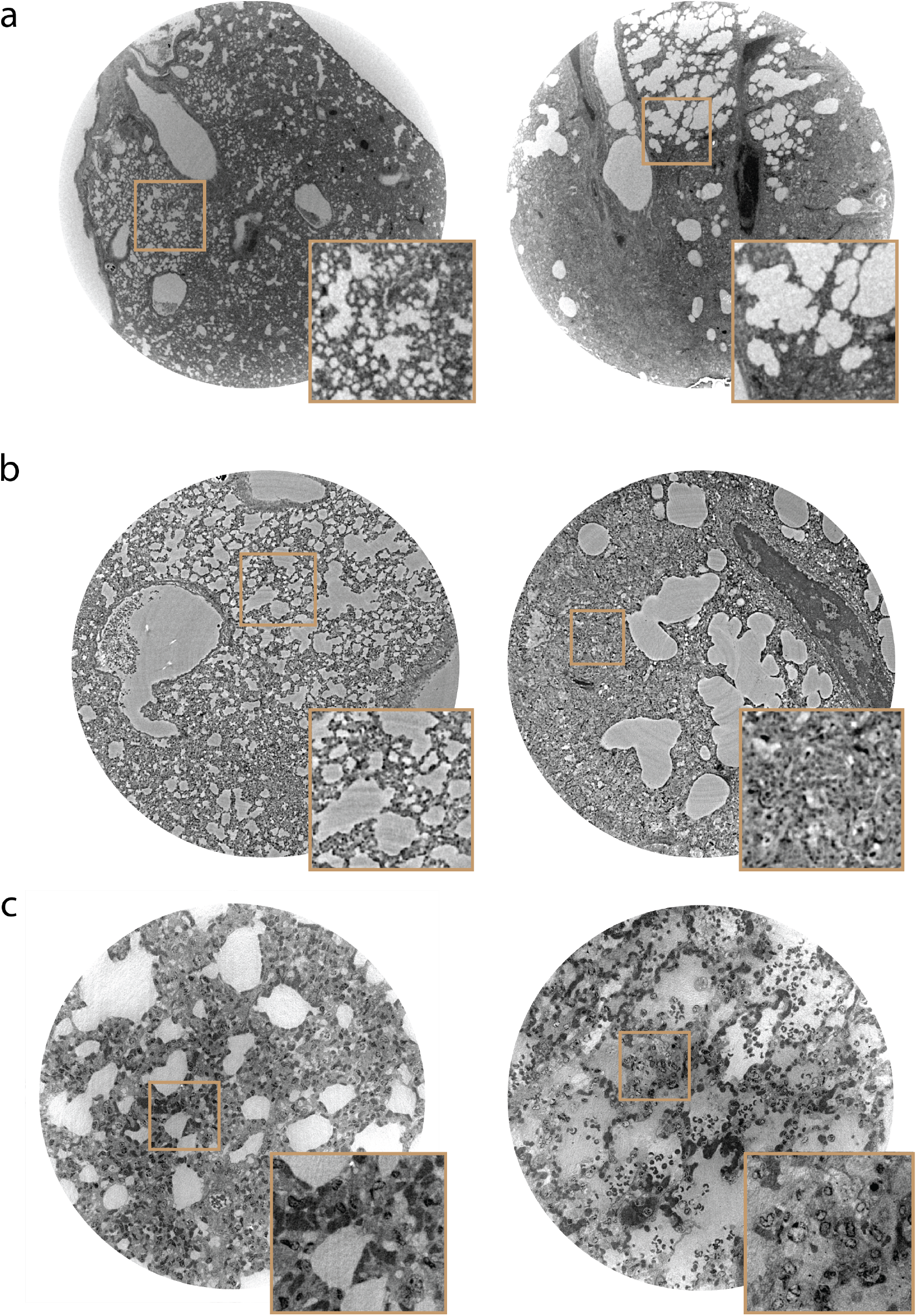
Comparison of imaging setups for uninfected control sample (left) and positive control (right): **a** Selected slices from in-house recorded tomograms. **b** Comparison of two slices recorded at the coherent imaging beamline at DESY (P10, PETRA III) with an effective pixel size of 650 nm and a FOV of 1.3 mm. **c** Slices from an uninfected (left) and positive control (right) hamster, recorded at the ID16a beamline (ESRF, Grenoble) with an effective pixel size of 90 nm and a FOV of 0.29 mm.

To achieve semi-automatic data acquisition and processing, up to 10 samples were stacked on top of each other inside a Kapton tube, and scanned as part of a single measurement run, controlled by a corresponding script. Subsequently, six different sample holders, each with up to 15 samples, were scanned under identical settings. Fig. 1**c** shows a photograph of the multi-sample holder in the beam, and Fig. 3**a** a sketch of the data recording scheme. After completion of the scans, the recorded volumes are reconstructed and processed by cropping a cube shaped volume of interest, followed by segmentation of air-filled compartments and tissue, as described next.

**Figure 3.**
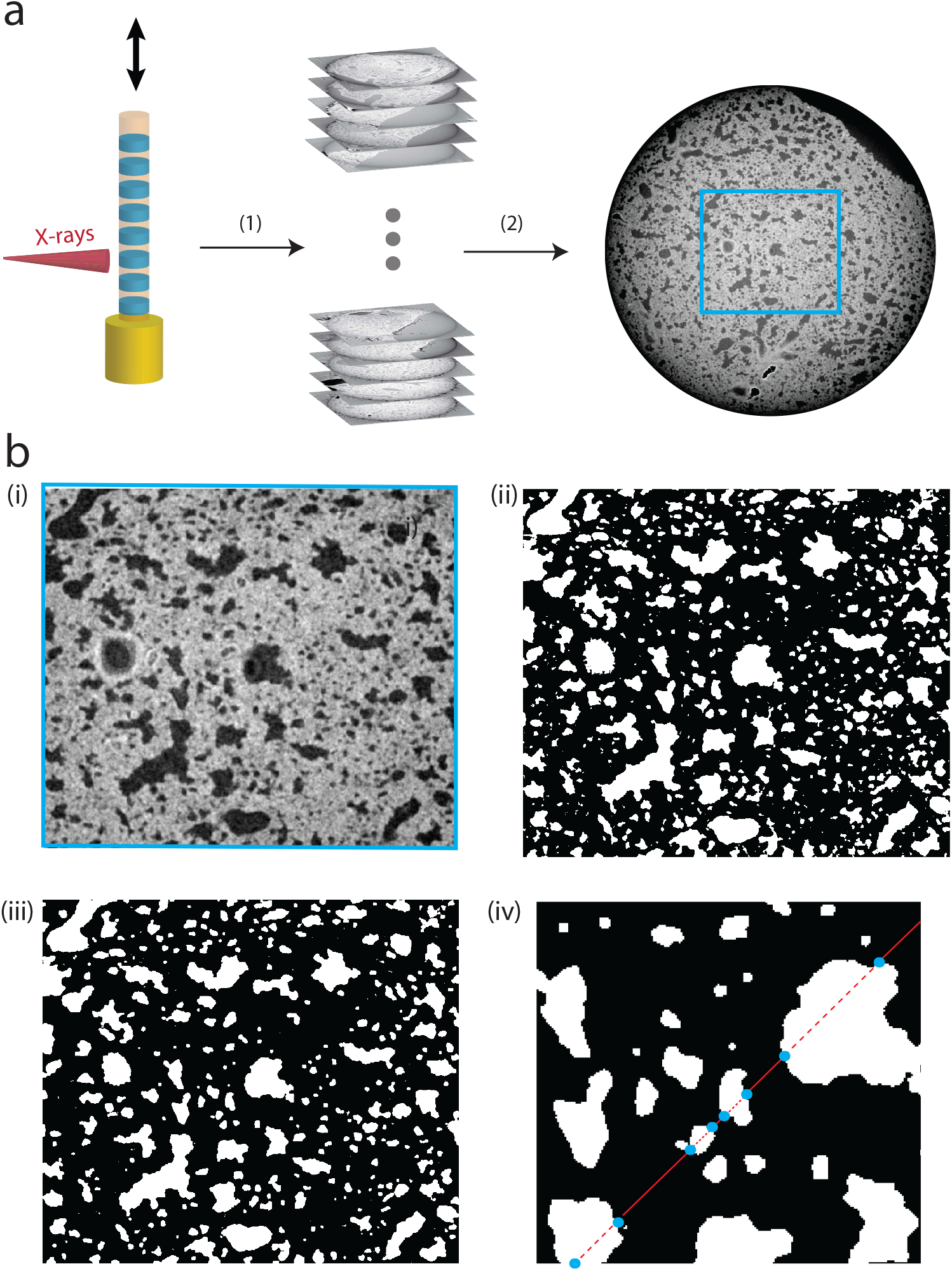
Demonstration of workflow for XPCT. **a** Illustration of data acquisition by using in-house XPCT. Paraffin-embedded sample biopsies are stacked onto each other in a Kapton tube and inserted into a Huber pin. By vertical translation all samples are scanned and the electron densities *ρ*_*x,y,z*_ are automatically reconstructed (1). Thereafter, a region of interest in the volume is selected (2). **b** (i) shows the selected region of interest, which is binned (ii) after finding a thresholding value (Otsu’s method [38]) and further processed by applying morphological operations (iii), namely a combination of erosion and dilation (opening/closing). Subsequently, chord lengths can be extracted by introducing a few thousand of randomly oriented lines to every slice of a binned volume. A line can pass through alveolar septae (phase 0, black) or alveolar lumen (phase 1, white) and is accordingly divided (iv). The length is determined by the distance between the intersection points (blue dots).

### Segmentation of Tissue and Determination of the Chord Length Distribution

Fig.3 demonstrates the successful implementation of image segmentation for later chord length analysis. Since the image contrast is sufficiently high with reduced level of artefacts, a computationally inexpensive and straightforward segmentation based on thresholding the gray values can be applied, i.e. Otsu’s thresholding method [38]. Note that an empirical offset was added to the threshold parameter, as controlled by visual inspection. This offset was kept constant for all samples. In combination with the subsequent 3D morphological operations (opening: 3 pixel diameter; closing: 1 pixel diameter sized spheres), a sufficiently accurate segmentation result is obtained for further processing steps. Fig. 3**b** illustrates the procedure step-by-step. With the volume mask at hand, the chord length distribution (CLD) can then be computed in a straightforward matter, as described in Sec. 2.

Figure 4 presents the CLD resulting from the image processing described above. Note that the normalized CLD represents the probability density function (PDF) for finding a chord of length *L*_*c*_ among all chords in the volume. In (a), this is shown for all hamsters, as a waterfall plot, with color indicating the CDL value, both for alveolar septae (phase 0) and the lumen (phase 1) In the lumen (phase 1), short chord lengths are significantly more prominent as expected with a peak at around 50 px, representing the mode of the distribution. Compared to the lumen, the CLD of the septae is significantly broader with a pronounced tail towards high *L*_*c*_. In (b) the PDFs are shown for hamsters of the (positive and negative) control groups only, enabling a better visual inspection than in the waterfall plot. In the data, one notes a tendency in the positive controls (red curves) to exhibit flatter and more extended distributions, compared to the negative controls (blue curves). The differences reflect a generally larger propensity of septae with a moderate thickness and a broadened and flattened distribution in the sick hamster lungs. Reciprocally, this may also imply a possible shrinkage or destruction of a larger fraction of alveolar spaces in infected lungs. However, one hamster (H59, black curve) appears to be a clear outlier and visual inspection of the dataset showed strong artifacts due to preparation, notably cracks in the paraffin. It was therefore discarded and excluded from further processing. Further, two outliers (H56, positive) and (H52, negative) exhibits curves closer to the respective opposite class. Note, that such outliers are not too surprising based on the biological variability as well as possible involuntary infection or wanted but failed infection. In the next, towards further analysis, each PDF is re-weighted by *L*_*c*_ to avoid that small chords have too much impact in the subsequent LOT analysis. In other words, larger chords corresponding to the right tails of the CLD are given more weight, facilitating the identification of characteristic differences between the different hamster lungs. The reweighted PDFs are again normalized, and on these reweighted PDFs LOT will be applied further below.

**Figure 4.**
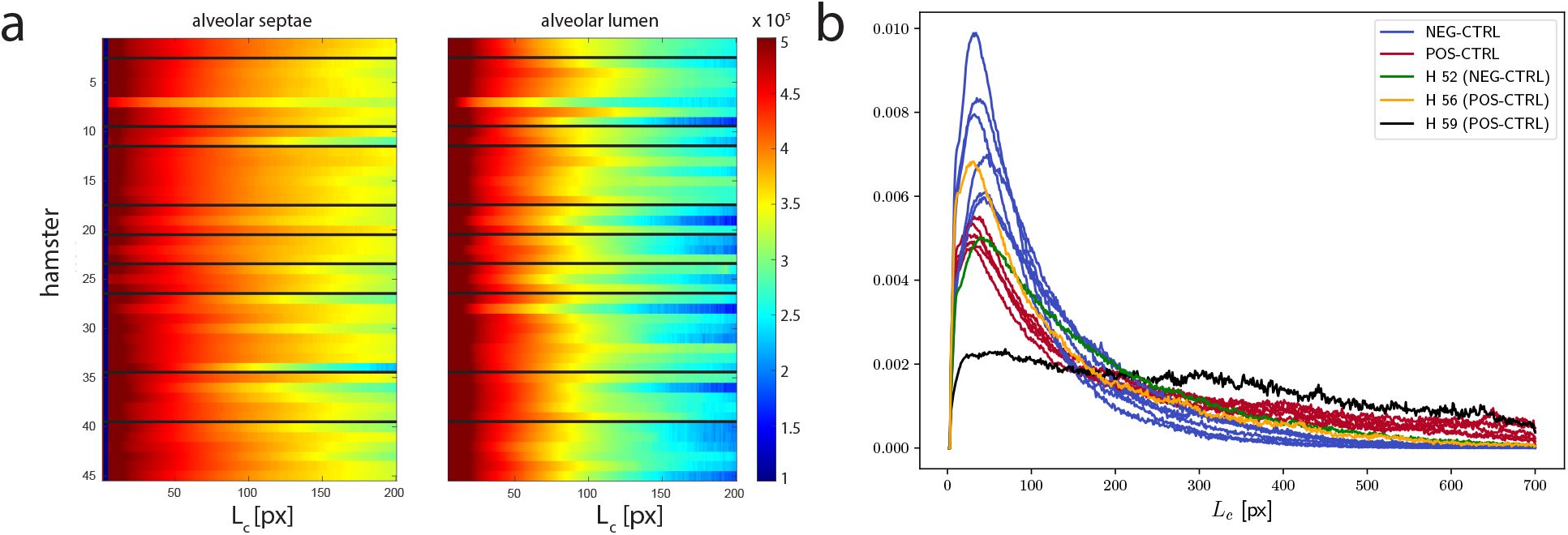
**a** Chord length distribution (CLD) for alveolar septae (left) and lumen (right), shown for all hamsters, including control groups and drug treated animals. The normalized CLD corresponds to the (unweighted) probability distribution function (PDFs) of finding a cord length,in the alveolar septae or lumen, respectively. **b** The PDFs for the septae data of the control groups (negative in blue, positive in red). Outliers are shown in green (H52, negative), orange (H56, positive), and black (H59). For the case of H59, inspection of the sample showed pronounced cracks in the paraffin, i.e. an artifact of sample preparation.

### Classification by Optimal Transport

In the following, the OT analysis based on inverse CDFs is applied to the post-processed hamster lung data, or more precisely to the re-weighted CLD of phase 0 (septae). We represent each sample by its inverse CDF and then apply PCA for dimensionality reduction.

### Control group

We began by analyzing the positive and negative control groups (POS-CTRL & NEG-CTRL) that should contain information on the changes in lung morphology associated with the pathological state, unaffected by drugs. The first two PCA components capture 94.5% and 4.9% of the dataset variance, indicating a clear low-dimensional structure. In the embedding Hamster H59 is separated by several standard deviations from the rest of the dataset, consistent with its very atypical chord length distribution (see Fig. 2 (b)), and indeed upon visual inspection of the 3D image, one can see a crack in the paraffin in the corresponding selected region in the scan. We therefore remove H59 as a clear outlier and continue the analysis without this sample. PCA now yields 97.8% and 1.5% of variance captured by the first two components. Fig. 5 (a) shows the embedded samples represented by their coordinates with respect to the two first PCA components. We observe that the coordinate along the first PCA component (PCA1) serves almost as a perfect classifier for the labels of the samples. With the exception of samples H52 and H56 all pathological samples have positive PCA1, all uninfected samples have a negative PCA1. As can be observed on Fig. 4 (b), samples H52 and H56 indeed appear to be more similar to the CLDs of the opposite class.

**Figure 5.**
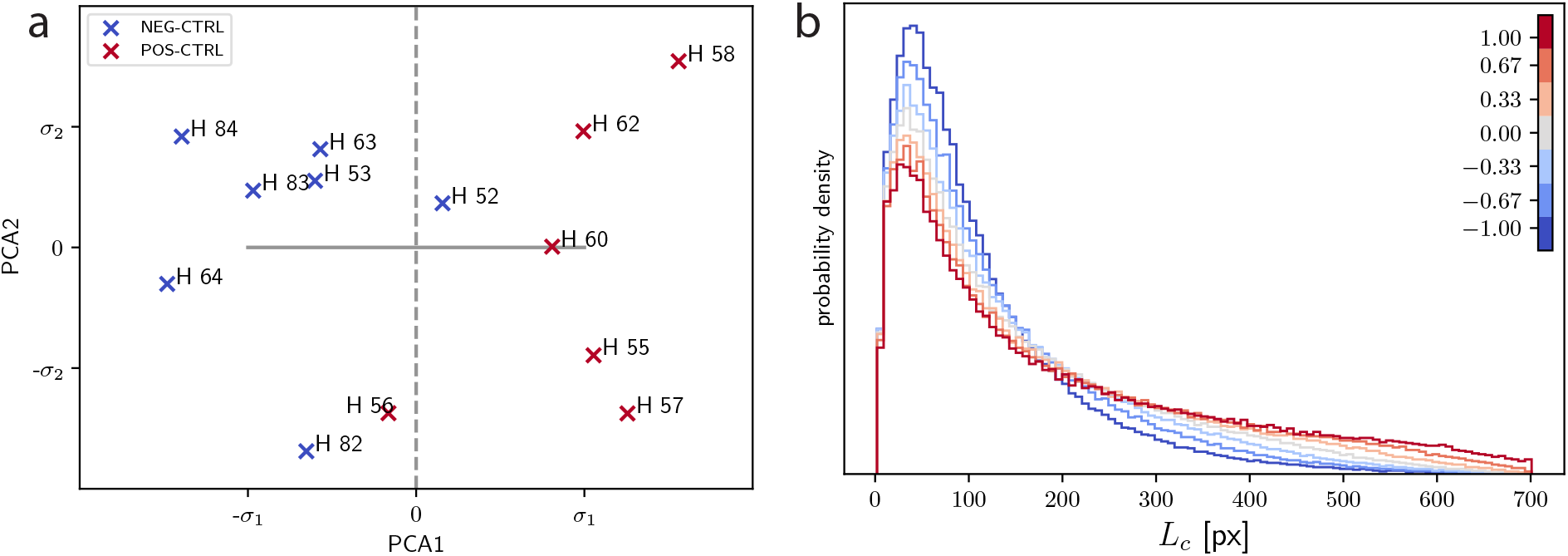
**a** Embedding of the control group into the plane spanned by the first two PCA components. Pathological (positive controls) hamsters are drawn in red while negative controls are in blue. Axes are scaled by the standard deviations *σ*_1_ and *σ*_2_ of the samples along each PCA component. The two subgroups are almost perfectly separated by their coordinate along the first PCA component (PCA1), as emphasized by the grey dashed line. **b** Evolution of the chord length distribution when moving from the origin along PCA1 by one standard deviation in both directions (represented by horizontal solid grey line in **a**).

A simple linear support vector machine (SVM) [50] applied to the embedding would likely be able to perfectly separate the samples by a straight line. But the orientation of this hyperplane will depend strongly on the small number of samples near the interface between the two classes and thus will be sensitive to noise. We will therefore use PCA1 as a more robust classifier for further analysis.

As mentioned above, there exists an inverse map from the two-dimensional PCA embedding via inverse CDFs to PDFs and therefore we can visualize hypothetical PDFs corresponding to a movement along the first PCA component by one standard deviation in both directions. This is shown in Fig. 5 (b). We observe moving into the pathological direction corresponds to a relative reduction of short chords. This is in line with the observation that SARS-CoV-2 infection is associated with thickening of septae. In the following, we can embed the other samples into the same PCA basis and then use PCA1 as an indicator for the strength of the pathological lung affection.

### Drug trial data

Now we examine the samples from the drug trial groups. First, we project them onto the first PCA component obtained from the control samples, to obtain their PCA1 values. Figure 6 shows these values of all drug samples clustered by groups, with means and standard deviations of each group. Drug groups 1 and 2 have consistently a relatively high PCA1 value (corresponding to strong pathology), whereas groups 3, 4, and 5 have lower values. While the low number of samples per drug candidate does of course not allow for a statistically definite estimation of the drugs efficiency, the relative consistency of the values suggests that candidates 3 to 5 are more promising for further development and evaluation than 1 and 2.

**Figure 6.**
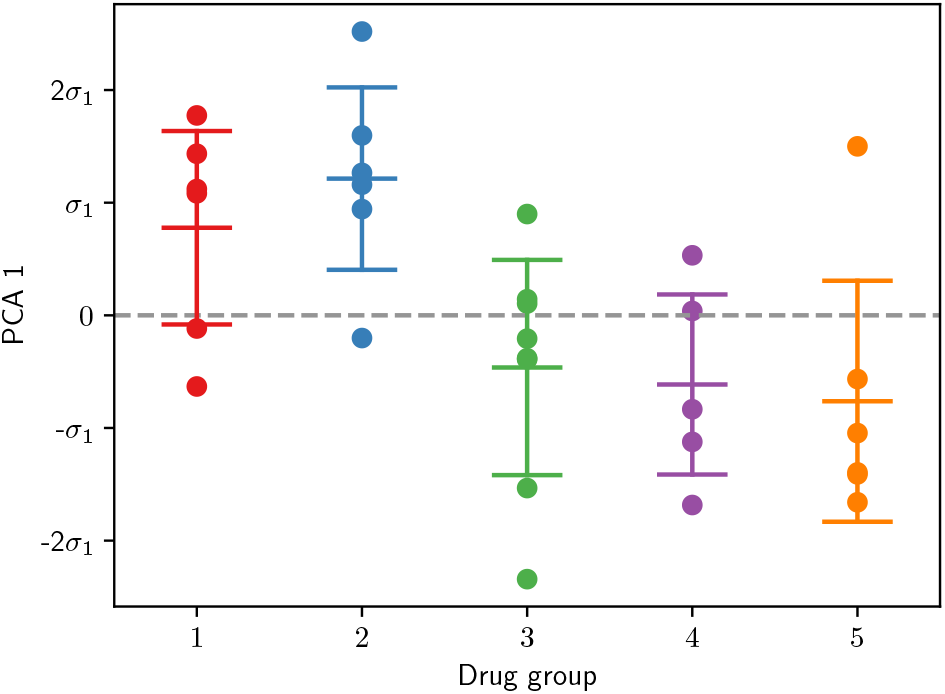
PCA1 values (in units of the standard deviation *σ*_1_ along the first principal component) of all five drug trials groups with mean and intervals of one standard deviation of each group indicated by lines. The dashed line indicates PCA1 = 0 for emphasis.

## 4 Discussion

We first discuss the analysis of the control group. This allowed us to attribute labels such as ‘healthy’ (uninfected/negative) and ‘sick’ (infected/positive) to the observed changes in the chord length distribution (CLD). We find that positive hamsters have a lower propensity for small chords (small alveolar spaces) and larger propensity for large chords, associated with larger empty entities, such as the non-physiological appearing fused spaces observed in the lung of the positive hamster shown on the left of Fig. 2. Note that while ‘healthy’ (uninfected/negative) and ‘sick’ (infected/positive) are dichotomous (binary) labels, the OT analysis and subsequent embedding equips us with a continuous score. We stress again that by comparing histograms with OT takes into account their full information (instead of only considering a few features such as their mean, median, variance, and similar) and is more geometrically robust than the L^2^-norm or the Kullback–Leibler divergence.

On this high-dimensional representation we then use PCA to extract a low-dimensional interpretable embedding. For the control data, more than 97 percent of the variance is captured by the first principal component (PCA1 axis in Fig.5). In other words, the hamsters of the control group are very well aligned along one direction in the high-dimensional space. In addition, this direction accounts almost perfectly for the prescribed labels. This is also a strong result in terms of the veterinary and imaging methodology. In fact, we would have expected more confounding variables due to the gitter in infection and recovery times of the small animals, as well as in the later sample preparation, imaging, and processing workflow. However, since the procedures were kept identical for all samples, it seems that the confounders were rather well controlled. For XPCS in particular, consistent segmentation is only warranted when the resolution, contrast, level of artifacts (illumination, raw data corrections, reconstruction errors) do not vary from sample to sample. While this can be difficult to achieve for high end synchrotron studies of highest resolution, we have here used a rather robust setting with high reproducibility at a time-stationary laboratory instrument.

Next, we discuss the drug candidates, i.e. the five drug groups which are classified with respect to the first PCA direction inferred from the control groups. A negative score indicates successful treatment as the lung recovered from infection, evidenced by the CLD regaining its physiological functional shape. A positive score means that the corresponding animals maintained the pathological morphology at the time of sacrifice. Here, we can make the following observations: while hamsters treated by drug group 1 and drug group 2 predominantly remain positive, hamsters treated by drug group 3, 4, and 5 have a negative (‘healthy’) average score, and a majority of hamsters on the negative side. Note that the drug compound and pharmaceutical background was not known to most co-authors, and in particular not to those co-authors involved in imaging and statistical testing. In fact, due to proprietary research, the information on the drugs cannot yet be disclosed, while the result of the present work is of course available to the companies involved. Note that for the methodological scope of this research it is not significant to know which drugs is which, and this ignorance even presents an advantage in view of a blinded and outcome independent investigation.

As main conclusions, we can note down:

- XPCT-based histopathology is compatible with pre-clinical studies, in view of the required *N*.
- XPCT in a laboratory setting and using a commercial instrument achieves sufficient image quality with reasonable loss compared to high-end synchrotron endstations.
- The chord length distribution (CLD) is a suitable morphometric descriptor for 3D lung tissue.
- Optimal transport (OT) can be efficiently implemented and can identify and describe the observed changes in the data.
- The transition from ‘sick’ to ‘healthy’ can be continuously visualized with respect to the CLD.
- The automated OT/PCA workflow can replace the operator-dependent lung affection score (LAS).

Finally, how to view the present OT/PCA-based approach in the wider field of machine learning? Compared to convolutional neural networks (CNN), the main advantage is that it is fully mathematically grounded, offers traceability, requires very few parameters to be set (such as the number of principal components to retain), and hence is explainable. At the same time, more work needs to be performed in view of quantitative statistical error margins. Owing to the one-dimensional distribution of the present data, the inverse cumulative density function transformation can be used, which makes it particularly accessible. The presented case is hence also ideally suited for the education of non-mathematicians. To this end, all relevant scripts, toolboxes and data are made available, along with sample documentation.

In the future, the assessment by trained physicians, radiologists, pathologists or veterinarians can be complemented by the automated workflow presented here. OT analysis in particular, but also the XPCT imaging workflow itself, could provide pre-clinical research and clinical practice with an augmented capability. In addition to reducing the workload of research or medical staff, automated assessment can improve the quality of research or diagnosis. For the case of drug development, throughput, sample volume, and data dimensionality could be much further increased. For early pre-clinical trials in particular, with low sample numbers the workflow could thus provide a valuable, robust analysis method with a very modest number of parameters. With a scan time of presently a little more than two hours per sample, studies of large animal cohorts can then also be accomplished in an acceptable amount of time. The subsequent data analysis can be implemented in the automatized workflow, minimizing the need for human and computational resources.

## Ethics statement

Hamster experiments were carried out according to the German Regulations for Animal Welfare after obtaining the necessary approval from the authorized ethics committee of the State Office of Agriculture, Food Safety and Fishery in Mecklenburg – Western Pomerania (LALLF MV) under permission number 7221.3-1-049/20 and approval of the commissioner for animal welfare at the Friedrich-Loeffler-Institute (FLI), representing the Institutional Animal Care and Use Committee (IACUC). The study is reported in accordance with ARRIVE guidelines.

## Data Availability

Raw data generated at ESRF/DESY will be released and made public two years after the beamtime. All treated datasets are available from the corresponding author on request. Exemplary data that support the findings and code used for data analysis will be openly available in GRO.data upon publication.

## Author Contributions Statement

TS, JR conceived the experiments. ABB and CB conducted the animal experiments and sample preparation at FLI. JR, TS conducted the synchrotron and in-house experiments. JR performed data processing, image reconstruction and visualization. BS, CS and SS provided the OT-framework and conducted the optimal transport analysis. JR, CS, SS, TS and BS interpreted the results. TS, BS, CS and JR wrote the manuscript. All authors reviewed the manuscript.

## Acknowledgements

We thank Markus Osterhoff, Michael Sprung and Fabian Westermeier for their continuous support at the instrument GINIX/P10 (PETRA III, DESY), where the reference scans of Fig.2 **b** were recorded, and Peter Cloetens for support as local contact during beamtime LS2980 at ID16a (ESRF), where the reference scan of Fig.2 **c** was recorded. The work was funded by the Deutsche Forschungsgemeinschaft (DFG, German Research Foundation) – Project-ID 432680300 – SFB 1456 and EXC 2067/1-390729940. We also acknowledge assistance in visualization with NVIDIA IndeX (NVIDIA Corporation, USA).

## Funding

Funded by the Deutsche Forschungsgemeinschaft (DFG, German Research Foundation) – Project-ID 432680300 – SFB 1456 and EXC 2067/1-390729940.

## Footnotes

1 https://pypi.org/project/LinOT/

## Notes

### Competing Interest Statement

The authors have declared no competing interest.

## References

1. Töpperwien, M. 3d virtual histology of neuronal tissue by propagation-based x-ray phase-contrast tomography. Göttingen Series in x-ray Physics (Göttingen University Press, Göttingen, 2018).

2. Töpperwien, M., van der Meer, F., Stadelmann, C. & Salditt, T. Three-dimensional virtual histology of human cerebellum by X-ray phase-contrast tomography. Proc. Natl. Acad. Sci. United States Am. 115, 6940–6945, DOI: 10.1073/pnas.1801678115 (2018).

3. Reichardt, M. et al. 3D virtual histopathology of cardiac tissue from Covid-19 patients based on phase-contrast X-ray tomography. eLife 10, e71359. DOI: 10.7554/eLife.71359 (2021). Publisher: eLife Sciences Publications, Ltd.

4. Parsons, D. W. et al. High-resolution visualization of airspace structures in intact mice via synchrotron phasecontrast X-ray imaging (PCXI). J. Anat. 213, 217–227, DOI: 10.1111/j.1469-7580.2008.00950.x (2008). _eprint: https://onlinelibrary.wiley.com/doi/pdf/10.1111/j.1469-7580.2008.00950.x.

5. O’Connell, D. W. et al. Accurate measures of changes in regional lung air volumes from chest x-rays of small animals. Phys. Medicine & Biol. 67, 205002, DOI: 10.1088/1361-6560/ac934d (2022).

6. Borisova, E. et al. Micrometer-resolution X-ray tomographic full-volume reconstruction of an intact post-mortem juvenile rat lung. Histochem. Cell Biol. 155, 215–226, DOI: 10.1007/s00418-020-01868-8 (2021).

7. Stahr, C. S. et al. Quantification of heterogeneity in lung disease with image-based pulmonary function testing. Sci. Reports 6, 29438, DOI: 10.1038/srep29438 (2016). Number: 1 Publisher: Nature Publishing Group.

8. Leong, A. F. T. et al. Real-time measurement of alveolar size and population using phase contrast x-ray imaging. Biomed. Opt. Express 5, 4024–4038, DOI: 10.1364/BOE.5.004024 (2014). Publisher: Optica Publishing Group.

9. Morgan, K. S. et al. Methods for dynamic synchrotron X-ray respiratory imaging in live animals. J. Synchrotron Radiat. 27, 164–175, DOI: 10.1107/S1600577519014863 (2020).

10. Bayat, S., Porra, L., Suortti, P. & Thomlinson, W. Functional lung imaging with synchrotron radiation: Methods and preclinical applications. Phys. Medica: Eur. J. Med. Phys. 79, 22–35, DOI: 10.1016/j.ejmp.2020.10.001 (2020). Publisher: Elsevier.

11. Bayat, S., Cercos, J., Fardin, L., Perchiazzi, G. & Bravin, A. Pulmonary vascular biomechanics imaged with synchrotron phase contrast microtomography in live rats. Eur. Respir. J. 60, DOI: 10.1183/13993003.congress-2022.1741 (2022). Publisher: European Respiratory Society Section: 01.05 - Clinical respiratory physiology, exercise and functional imaging.

12. Shaker, K., Häggmark, I., Reichmann, J., Arsenian-Henriksson, M. & Hertz, H. M. Phase-contrast X-ray tomography resolves the terminal bronchioles in free-breathing mice. Commun. Phys. 4, 259, DOI: 10.1038/s42005-021-00760-8 (2021).

13. Bayat, S., Fardin, L., Cercos-Pita, J. L., Perchiazzi, G. & Bravin, A. Imaging Regional Lung Structure and Function in Small Animals Using Synchrotron Radiation Phase-Contrast and K-Edge Subtraction Computed Tomography. Front. Physiol. 13 (2022).

14. Eckermann, M. et al. 3D virtual pathohistology of lung tissue from Covid-19 patients based on phase contrast X-ray tomography. eLife 9, e60408, DOI: 10.7554/eLife.60408 (2020). Publisher: eLife Sciences Publications, Ltd.

15. van Griethuysen, J. J. et al. Computational Radiomics System to Decode the Radiographic Phenotype. Cancer research 77, e104–e107, DOI: 10.1158/0008-5472.CAN-17-0339 (2017).

16. Lovric, G. et al. Automated computer-assisted quantitative analysis of intact murine lungs at the alveolar scale. PLOS ONE 12, e0183979, DOI: 10.1371/journal.pone.0183979 (2017).

17. Kumar, V. et al. Radiomics: the process and the challenges. Magn. Reson. Imaging 30, 1234–1248, DOI: 10.1016/j.mri.2012.06.010 (2012).

18. Lambin, P. et al. Radiomics: Extracting more information from medical images using advanced feature analysis. Eur. J. Cancer 48, 441–446, DOI: 10.1016/j.ejca.2011.11.036 (2012).

19. Peyré, G. & Cuturi, M. Computational Optimal Transport. arXiv:1803.00567 [stat] (2020). ArXiv: 1803.00567.

20. Foxley, S. et al. Multi-modal imaging of a single mouse brain over five orders of magnitude of resolution. NeuroImage 238, 118250, DOI: 10.1016/j.neuroimage.2021.118250 (2021).

21. Haker, S., Zhu, L., Tannenbaum, A. & Angenent, S. Optimal Mass Transport for Registration and Warping. Int. J. Comput. Vis. 60, 225–240, DOI: 10.1023/B:VISI.0000036836.66311.97 (2004).

22. Rabin, J. & Papadakis, N. Convex Color Image Segmentation with Optimal Transport Distances (2015). ArXiv:1503.01986 [cs].

23. Guo, Y., Wang, X., Li, C. & Ying, S. Domain adaptive semantic segmentation by optimal transport. Fundamental Res. DOI: 10.1016/j.fmre.2023.06.006 (2023).

24. Schmitzer, B. & Schnörr, C. Object Segmentation by Shape Matching with Wasserstein Modes. In Heyden, A., Kahl, F., Olsson, C., Oskarsson, M. & Tai, X.-C. (eds.) Energy Minimization Methods in Computer Vision and Pattern Recognition, Lecture Notes in Computer Science, 123–136, DOI: 10.1007/978-3-642-40395-8_10 (Springer, Berlin, Heidelberg, 2013).

25. Crook, O. M. et al. A Linear Transportation $\mathrm{L}^p$ Distance for Pattern Recognition (2020). ArXiv:2009.11262 [cs, math].

26. Courty, N., Flamary, R., Tuia, D. & Corpetti, T. Optimal transport for data fusion in remote sensing. In 2016 IEEE International Geoscience and Remote Sensing Symposium (IGARSS), 3571–3574, DOI: 10.1109/IGARSS.2016.7729925 (2016). ISSN: 2153-7003.

27. Muñoz-Fontela, C. et al. Animal models for COVID-19. Nature 586, 509–515, DOI: 10.1038/s41586-020-2787-6 (2020). Number: 7830 Publisher: Nature Publishing Group.

28. Blaurock, C. et al. Compellingly high SARS-CoV-2 susceptibility of Golden Syrian hamsters suggests multiple zoonotic infections of pet hamsters during the COVID-19 pandemic. Sci. Reports 12, 15069, DOI: 10.1038/s41598-022-19222-4 (2022). Number: 1 Publisher: Nature Publishing Group.

29. Wölfel, R. et al. Virological assessment of hospitalized patients with COVID-2019. Nature 581, 465–469, DOI: 10.1038/s41586-020-2196-x (2020). Number: 7809 Publisher: Nature Publishing Group.

30. Reichmann, J. et al. Human lung virtual histology by multi-scale x-ray phase-contrast computed tomography. Phys. Medicine & Biol. 68, 115014, DOI: 10.1088/1361-6560/acd48d (2023). Publisher: IOP Publishing.

31. Frohn, J. et al. 3D virtual histology of human pancreatic tissue by multiscale phase-contrast X-ray tomography. J. Synchrotron Radiat. 27, 1707–1719, DOI: 10.1107/S1600577520011327 (2020).

32. Cloetens, P. et al. Holotomography: Quantitative phase tomography with micrometer resolution using hard synchrotron radiation x rays. Appl. Phys. Lett. 75, 2912–2914, DOI: 10.1063/1.125225 (1999).

33. Lohse, L. et al. A phase-retrieval toolbox for X-ray holography and tomography. J. Synchrotron Radiat. 27, DOI: 10.1107/S1600577520002398 (2020).

34. Yu, B. et al. Evaluation of phase retrieval approaches in magnified X-ray phase nano computerized tomography applied to bone tissue. Opt. Express 26, 11110–11124, DOI: 10.1364/OE.26.011110 (2018). Publisher: Optica Publishing Group.

35. van Aarle, W. et al. The ASTRA Toolbox: A platform for advanced algorithm development in electron tomography. Ultramicroscopy 157, 35–47, DOI: 10.1016/j.ultramic.2015.05.002 (2015).

36. Mirone, A., Brun, E., Gouillart, E., Tafforeau, P. & Kieffer, J. The PyHST2 hybrid distributed code for high speed tomographic reconstruction with iterative reconstruction and a priori knowledge capabilities. Nucl. Instruments Methods Phys. Res. Sect. B: Beam Interactions with Mater. Atoms 324, 41–48, DOI: 10.1016/j.nimb.2013.09.030 (2014).

37. Knudsen, L., Weibel, E. R., Gundersen, H. J. G., Weinstein, F. V. & Ochs, M. Assessment of air space size characteristics by intercept (chord) measurement: an accurate and efficient stereological approach. J. Appl. Physiol. 108, 412–421, DOI: 10.1152/japplphysiol.01100.2009 (2010). Publisher: American Physiological Society.

38. Otsu, N. A Threshold Selection Method from Gray-Level Histograms. IEEE Transactions on Syst. Man, Cybern. 9, 62–66, DOI: 10.1109/TSMC.1979.4310076 (1979). Conference Name: IEEE Transactions on Systems, Man, and Cybernetics.

39. Bresenham, J. E. Algorithm for computer control of a digital plotter. IBM Syst. J. 4, 25–30, DOI: 10.1147/sj.41.0025 (1965). Conference Name: IBM Systems Journal.

40. MacIver, M. R. Chord Length Distribution from Binary 2D Images (2023).

41. Chung, S.-Y., Sikora, P., Rucińska, T., Stephan, D. & Abd Elrahman, M. Comparison of the pore size distributions of concretes with different air-entraining admixture dosages using 2D and 3D imaging approaches. Mater. Charact. 162, 110182, DOI: 10.1016/j.matchar.2020.110182 (2020).

42. Turner, D. M., Niezgoda, S. R. & Kalidindi, S. R. Efficient computation of the angularly resolved chord length distributions and lineal path functions in large microstructure datasets. Model. Simul. Mater. Sci. Eng. 24, 075002, DOI: 10.1088/0965-0393/24/7/075002 (2016). Publisher: IOP Publishing.

43. Latypov, M. I. et al. Application of chord length distributions and principal component analysis for quantification and representation of diverse polycrystalline microstructures. Mater. Charact. 145, 671–685, DOI: 10.1016/j.matchar.2018.09.020 (2018).

44. Crowley, G. et al. Quantitative lung morphology: semi-automated measurement of mean linear intercept. BMC Pulm. Medicine 19, 206, DOI: 10.1186/s12890-019-0915-6 (2019).

45. Santambrogio, F. Optimal Transport for Applied Mathematicians: Calculus of Variations, PDEs, and Modeling (Birkhäuser, 2015). Google-Books-ID: UOHHCgAAQBAJ.

46. scipy.interpolate.interp1d — SciPy v1.11.2 Manual.

47. Park, S. & Thorpe, M. Representing and Learning High Dimensional Data with the Optimal Transport Map from a Probabilistic Viewpoint. In 2018 IEEE/CVF Conference on Computer Vision and Pattern Recognition, 7864–7872, DOI: 10.1109/CVPR.2018.00820 (IEEE, Salt Lake City, UT, 2018).

48. Wang, W., Slepčev, D., Basu, S., Ozolek, J. A. & Rohde, G. K. A Linear Optimal Transportation Framework for Quantifying and Visualizing Variations in Sets of Images. Int. J. Comput. Vis. 101, 254–269, DOI: 10.1007/s11263-012-0566-z (2013).

49. Frost, J. et al. 3d virtual histology reveals pathological alterations of cerebellar granule cells in multiple sclerosis. preprint, Pathology (2022). DOI: 10.1101/2022.10.07.22280811.

50. Cortes, C. & Vapnik, V. Support-vector networks. Mach. Learn. 20, 273–297, DOI: 10.1007/BF00994018 (1995).

