## Supplemental Document 1 for "Cytoarchitecture of SARS-CoV-2 infected hamster lungs by X-ray phase contrast tomography: imaging workflow and classification for drug testing"

We here present additional graphics and tables, detailing in particular data concerning the viral load measured after sample extraction by nasal swab, for different days post infection (dpi) (Tab. 1 and Fig. 1)). Fig. 2 illustrates the re-weighting procedure of the chord length, by an example plot, and Fig. 3 illustrates the relative change in chord length along the PCA direction, as well as a classification by support vector machine (SVM). Finally, Fig. 4 presents the correlation of the PCA1 component and the lung affectation score (LAS), both for (a) the control group, and (b) the drug-treated hamsters.

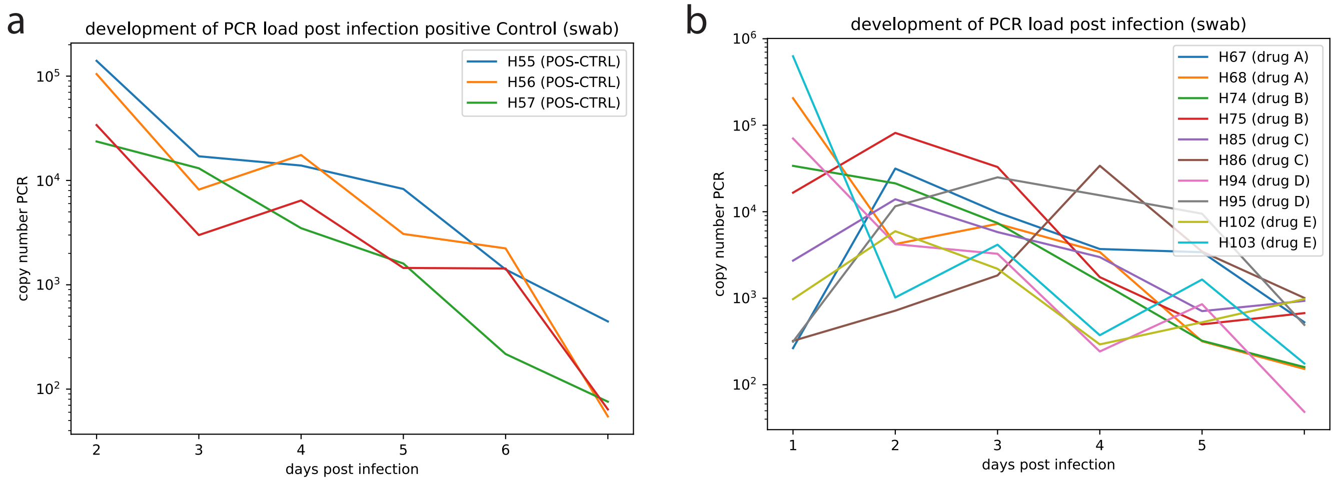


**Figure 1.** PCR viral load. **a** PCR copy number per $\mu l$ sample as a function of day post infection (dpi) in the positive control group, and **b** in the drug-treated hamsters (selected examples only, complete data see Tab. 1). Samples were collected by nasal wash.

| | 0 dpi cnPCR [$\mu l$] | 1 dpi cnPCR [$\mu l$] | 2 dpi cnPCR [$\mu l$] | 3 dpi cnPCR [$\mu l$] | 4 dpi cnPCR [$\mu l$] | 5 dpi cnPCR [$\mu l$] |
|---|---|---|---|---|---|---|
| H67 | 265 | 31570 | 9804 | 3707 | 3405 | 528 |
| H68 | 205584 | 4231 | 7250 | 3419 | 318 | 152 |
| H74 | 33918 | 21290 | 7397 | 1561 | 322 | 160 |
| H75 | 16612 | 81580 | 32907 | 1754 | 499 | 673 |
| H85 | 2721 | 13939 | 5820 | 2989 | 710 | 932 |
| H86 | 321 | 718 | 1837 | 34015 | 3464 | 1008 |
| H94 | 70295 | 4218 | 3266 | 242 | 854 | 49 |
| H95 | 314 | 11570 | 24988 | 15495 | 9487 | 492 |
| H102 | 977 | 5956 | 2189 | 293 | 527 | 979 |
| H103 | 627559 | 1021 | 4160 | 374 | 1647 | 176 |

**Table 1.** SARS-CoV-2 gene copy number per $\mu l$ sample exemplified on hamsters from different drug groups (plotted in Fig1c)

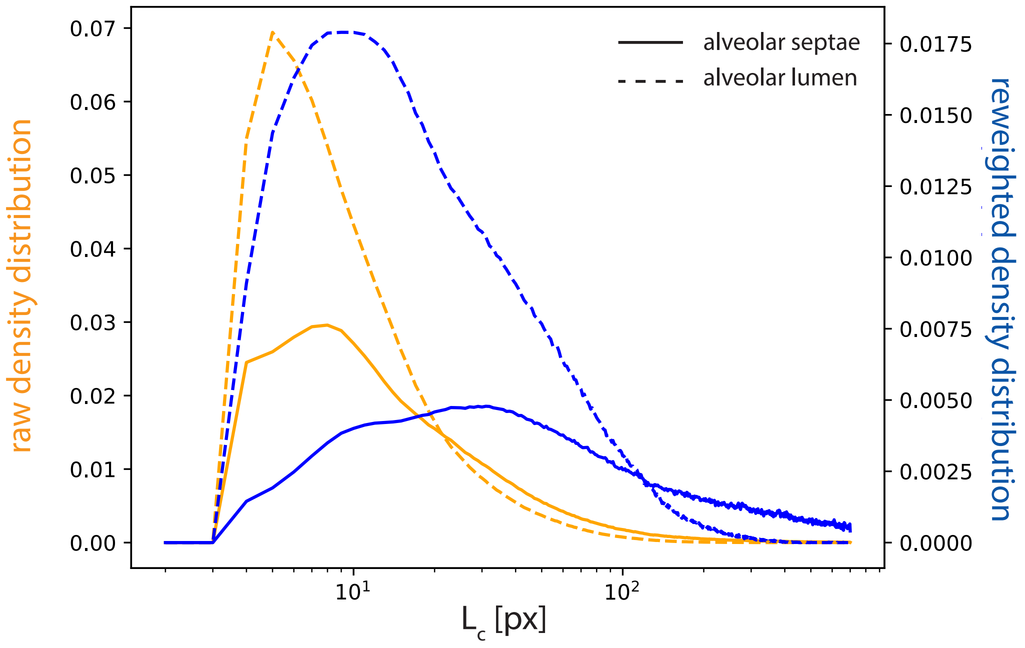


**Figure 2.** Illustration of reweighting. Probability distribution function (pdf) of finding chords of length $L_c$ in septae (solid lines) and alveolar lumen (dashed lines), for the unweighted (yellow) and $L_c$-weighted (blue) case. Curves are shown for an exemplary POS-CTRL hamster.

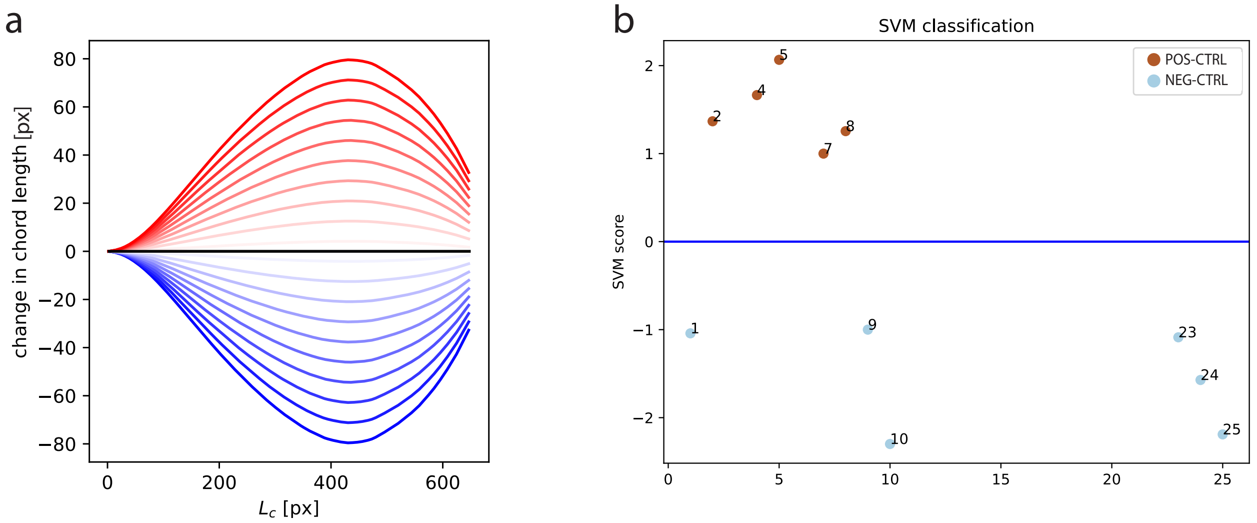


**Figure 3.** Relative change in CLD and SVM classification. **a** In addition to the frequency of the cords, i.e. the CLD, we can also compute the changes in chord length, when moving along PC1. The results quantify the thickening of the septae as one moves from negative (healthy) to positive (sick). **b** Distance from the hyperplane in the two-dimensional PCA space, for the control samples. The distance to the hyperplane (blue line) defines the SVM score.

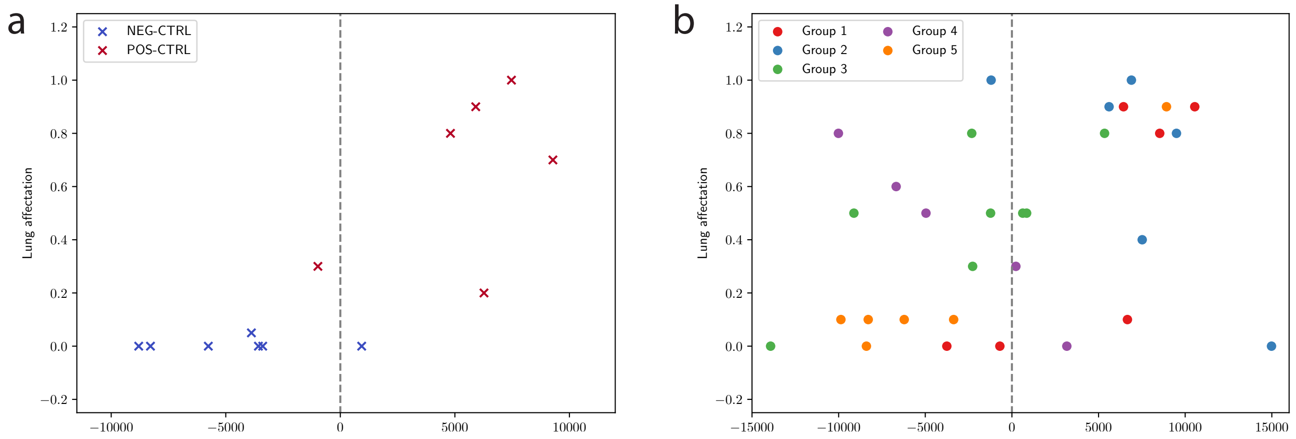


**Figure 4.** Representation of control (a) and drug (b) groups by their PCA 1 coordinate, plotted against their LA score. The separator at $PCA1 = 0$ is drawn as a dashed line. **a** Representation of negative (blue) and positive (red) control samples along PCA1 in the embedding space with respect to the attributed lung affectation score and the subgroups being separated by their first PCA coordinate (dashed gray line). **b** Illustration of drug groups along PCA1 in the embedding space with according to their lung affectation score.
